# Real time analysis of SARS-CoV-2 induced cytolysis reveals distinct variant-specific replication profiles

**DOI:** 10.1101/2023.03.28.534588

**Authors:** Sarah E Scheuermann, Kelly Goff, Lori A Rowe, Brandon J Beddingfield, Nicholas J Maness

**Affiliations:** Division of Microbiology, Tulane National Primate Research Center, Covington, LA 70433, USA; Department of Microbiology and Immunology, Tulane University School of Medicine, New Orleans, LA 70112, USA

## Abstract

The continuous evolution of new SARS-CoV-2 variants with enhanced immune evasion capacity suggests the entire population is and will continue to be potentially vulnerable to infection despite pre-existing immunity. The ability of each new variant to evade host humoral immunity is the focus of intense research across the globe. Each variant may also harbor unique replication capabilities relevant for disease and transmission. Here we demonstrate the utility of a new approach to assessing viral replication kinetics using Real Time Cell Analysis (RTCA). Virus induced cell death is measured in real time by the detection of electrical impedance through cell monolayers. Using this system, we quantified replication kinetics of five clinically important viral variants; USA WA1/2020 (an A1 ancestral lineage isolate), Delta, and Omicron subvariants BA.1, BA.4, and BA.5. We identified multiple kinetic measures that proved useful in variant replication comparisons including time (in hours) to the maximum rate of cell death at each log10 viral dilution and the slope at the maximum rate of cell death. We found that WA1/2020 and Delta were the most rapid but in distinct ways. While WA1/2020 induced cell death most rapidly after inoculation, Delta was slightly slower to reach cell death, it appeared to kill cells faster once cytotoxic effects began. Interestingly, BA.1, showed substantially reduced replication kinetics relative to all other variants. Together, these data show that real time analysis of cell death is a robust method to assess replicative capacity of any given SARS-CoV-2 variant rapidly and quantitatively, which may be useful in assessment of newly emerging variants.

## Introduction

The recently emerged coronavirus, SARS-CoV-2, has been responsible for the ongoing pandemic since 2019. Since its initial emergence, it has continuously evolved, fueled by immune pressure from vaccination and natural infection in the human population. From the early widely circulating strain, deemed WA1/2020 in the United States [1], variants have included the first D614G mutant [2,3], Alpha (B.1.1.7) [4,5], Beta (B.1.351) [6], Gamma (P.1) [7,8], Delta (B.1.617.2) [9,10], Lambda (C.37) [11], and Mu (B.1.621) [12]. Since it was first detected in November of 2021, the variant of concern termed Omicron has spawned multiple sub-lineages, many of which show substantial variation relative to the original Omicron variant from which they evolved [13,14]. New variants/sub-variants are routinely discovered, such as Omicron XBB.1.5 [15] and BQ.1.1 [16], due to recombination within hosts, immune evasion, and spread within populations.

New variants rapidly spread through populations and often become dominant before declining in frequency and being replaced by a new variant, often with even greater antibody escape capacity [9,10,14,17,18]. This has resulted in monoclonal antibody therapeutics and infection-derived antibodies becoming ineffective as the pandemic has continued, due to the highly specific nature of antibody binding [16,19-22]. Vaccines are affected as well, though often to a lesser degree [6,13,23-27]. Immunity generated by the widely used mRNA vaccines also shows decreased neutralizing capacity for variants relative to the ancestral variant, with serum from vaccinees having up to 3-fold decreased neutralization for the Delta variant [28] and far greater reductions in neutralization of omicron and its subvariants. Variants may evade cellular immunity as well [29] though this appears to be less widespread than evasion of humoral immunity.

In addition to immune escape, mutations in newly emerged variants may also impact viral infection, replication, and transmission. Mutations within the receptor binding domain (RBD) of the spike protein may modify binding and uptake of virus while spike mutations outside the RBD may alter aspects of replication in other ways favorable for intrahost viral dynamics. Altered replication dynamics may also impact disease severity and inter-host transmission [3,17,30]. In prior work, Omicron showed reduced viral replication kinetics in cell culture, potentially due to a reduced ability to antagonize the interferon response as compared to Delta [31], which may be associated with the reduced severity of disease associated with the Omicron variant. Mutations underlying these altered kinetics may lie outside of spike in structural or nonstructural proteins [32]. Altogether, these findings highlight the importance of robust characterization of viral kinetics in live, whole virus assays.

A technology utilized in the cancer biology space that has expanded into virology laboratories in recent years, termed Real-Time Cell Analysis (RTCA) using the Agilent xCELLigence eSight system, has the potential to rapidly assess viral replication kinetics and other important parameters. This technology is based upon real time measurements of electrical impedance of a cell monolayer [33-38]. The impedance is a correlate of monolayer integrity, with impedance falling as cells are destroyed, including as a result of lytic viral infection. This is reflected in a unitless value termed cell index, which can be monitored on a per-well basis over the course of multiple days and combined with visual monitoring of the monolayer using an integrated microscope and camera. We herein describe the use this technology as a platform for detailed examination of the *in vitro* kinetics of replication of multiple SARS-CoV-2 variants of concern. We show RTCA to be an ideal tool for viral characterization that can aid in elucidation of unique aspects of emerging variants during a rapidly evolving pandemic.

## Materials and Methods

### Virus and Cells

Multiple SARS-CoV-2 variants were used to infect Vero-TMPRSS2 cells (# JCRB1819, JCRB Cell Bank). Cells were cultured in Dulbecco’s modified Eagle’s medium (DMEM) supplemented with 10% fetal bovine serum (FBS) and 1% Anti-Anti additionally supplemented with 2% G418 Sulfate Solution.

The following viruses were received from BEI: NR-54001, icSARS-CoV-2-WT (WA1/2020); hCoV-19/USA/MD-HP20874/2021 WCCM (Omicron BA.1): NR-58620, hCoV-19/USA/COR-22-063113/2022 WCCM (Omicron BA.5); and NR-56806, hCoV-19/USA/MD-HP30386/2022 WCCM (Omicron BA.4); NR-55672, hCoV-19/USA/MD-HP05647/2021 (Delta B.1.617.2). Virus was propagated in Vero-TMPRSS2 cells to create stocks. Sequences of new stocks were confirmed by Ilumina sequencing as previously described. Genome assembly and variant analysis was performed using DRAGEN COVID Lineage pipeline as an Illumina BaseSpace App following standard protocol, except for a custom primer BED file containing the SWIFT primers [39].

### TCID_**50**_

Median Tissue Culture Infectious Dose (TCID_50_) was performed on each stock to quantify the amount of active, replication competent virus. Vero TMPRSS2 cells were plated in 48-well tissue culture treated plates to be subconfluent at time of assay. Cells were washed with serum free DMEM and 50uL of virus was allowed to adsorb onto the cells for 1 hour at 37°C and 5% CO2. After adsorption, cells were overlayed with DMEM containing 2% FBS and 1% Anti/Anti (#15240062, Thermo Scientific, USA). Plates were incubated for 7–10 days before being observed for cytopathic effect (CPE). Any CPE observed relative to control wells was considered positive and used to calculate TCID_50_ by the Reed and Muench method [40].

### Real Time Cell Analysis assay setup and data analysis

Vero TMPRSS2 cells were plated in 96-well tissue culture treated E-Plate VIEW plates (#300-601-020, Agilent) to be subconfluent at time of assay. Viral strains were each diluted with DMEM containing 2% FBS and 1% Anti/Anti to the same starting concentration of 1e5 TCID_50_ followed by 1:10 serial dilutions for a total of seven dilutions. Media was removed from the wells of the 96 well plate, and 100uL of virus samples were added. 100uL of 2% FBS 1% Anti/Anti DMEM were added to the negative control wells. The plates were then placed on the xCELLigence RTCA eSight impedance and imaging cradles (cradles 1-3). The plate layouts and experiment schedule were defined in the Esight software. Impedance measurements for each well were collected every 15 minutes and images for each well collected every 60 minutes over the course of 5 days.

Cell index values over time were graphed in the xCELLigence software. Graphs for all plates were then normalized at the same timepoint (11.76 hours) with the delta cell index function to add a constant to the cell index of each well. Area Under the Curve (AUC) for each replicate was calculated using the Area Under the Curve analysis function in the Prism software. The AUC baseline parameters were set based on the lowest delta cell index (impedance) value for each replicate. Kruskal-Wallis test was then performed to compare the total AUC of variants at each concentration.

Slope values over time were graphed in the xCELLigence software and normalized to the same timepoint (11.76 hours) prior to exporting data. The lowest slope (steepest downward slope) value of each replicate was identified as the “max slope”. The time at which each replicate reached the max slope was also recorded. Kruskal-Wallis test was then performed to compare the value and time of max slopes of each variant at each concentration.

## Results

### Cell index patterns differ between variants

Vero/TMPRSS2 cells were inoculated with multiple SARS-CoV-2 variants of concern to characterize the viral ability to infect and destroy the monolayer, by near-continuous monitoring of the monolayer impedance to electricity as a correlate of cellular death. Readings of the monolayer impedance were taken every fifteen minutes over the course of five days in order to generate impedance curves (Fig. S1). Various characteristics of these curves were analyzed, including time to maximum (max) slope, value of max slope, and area under the curve (AUC) of the impedance drop. These were performed using multiple dilutions of virus in order to characterize differences at a range of infectious doses. The specifics of these patterns and their differences by variant are captured in this manner.

### Slope characteristics differ between variants

The time taken to reach the max slope was the first metric used to assess replication kinetics of the different variants across a range of dilutions. In all dilutions, WA1/2020 reached max slope more quickly than all other variants, save for Delta at 1e4 and 1e3 TCID_50_, though the trend persisted with those comparisons as well. BA.1 was the slowest at reaching max slope across dilutions, though not all pairwise comparisons reached statistical significance. BA.4, BA.5 and Delta were not significantly different from each other across dilutions (Fig 1A).

**Fig. 1.**
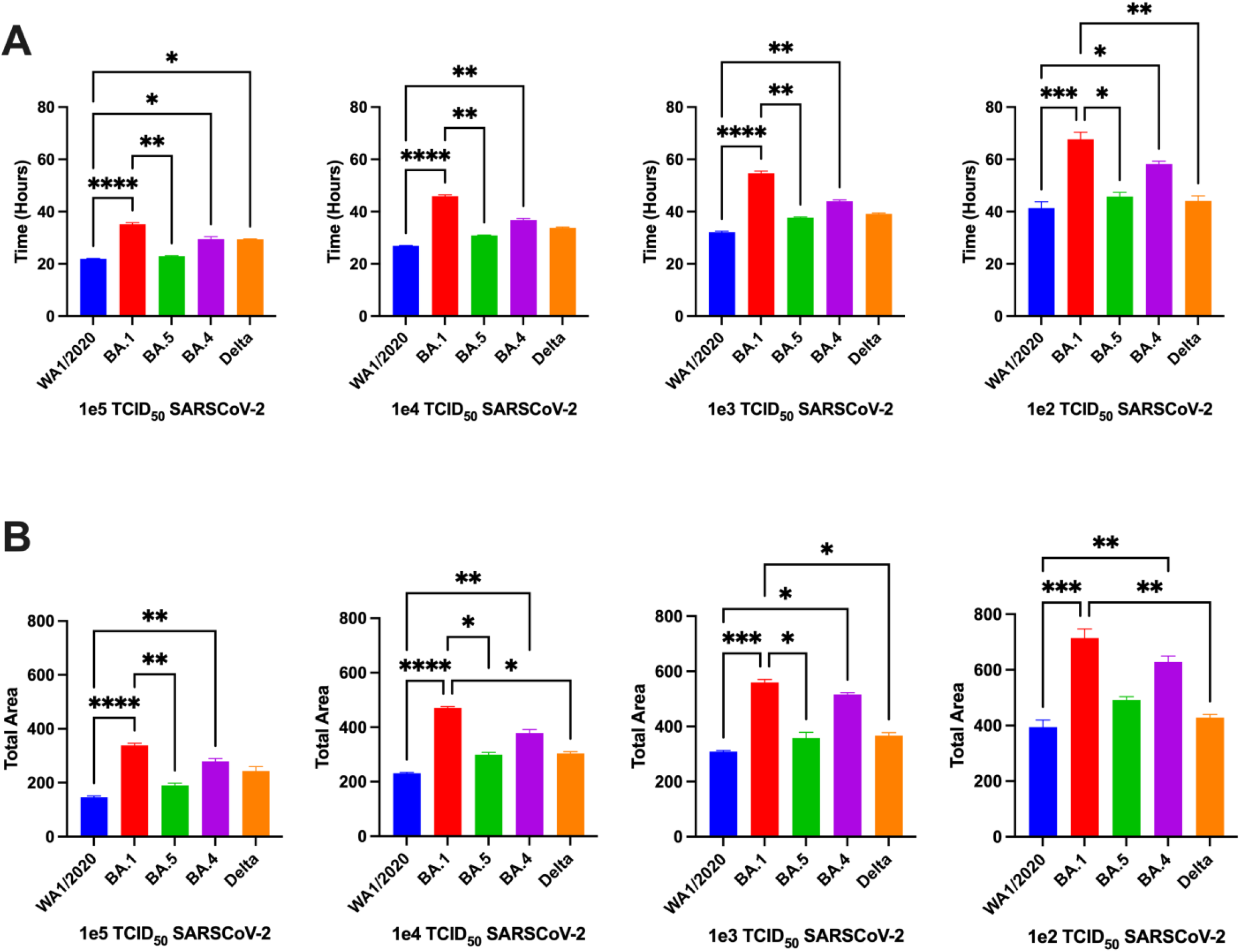
Kinetic Comparisons of Vero/TMPRSS2 cells inoculated with multiple SARS-CoV-2 Variants of Concern. (A) Time to reach max slope of inoculated cells and (B) Area under the curve of cell index. Data is represented as mean with SEM. Groups were compared via Kruskal-Wallis test. (*, p<0.05; **, p<0.01; ***, p<0.001; ****, p<0.0001).

The area under the cell index curve (AUC) was next analyzed for each variant and dilution. WA1/2020 had the lowest AUC across all dilutions, being significantly lower than BA.1, BA.4, and BA.5 at various dilutions, but never reaching significance compared to Delta. BA.1 had the highest AUC of the variants at all dilutions, being significantly higher than Delta at 3 of 4 dilutions and higher than WA1/2020 at all dilutions (Fig. 1B).

The absolute value of the slope generated by each variant was also assessed. A clear pattern emerged wherein Delta exhibited the steepest absolute slope while BA.1 exhibited the least steep slope (Figure 2A). Statistical analyses validated these observations. Delta generated the steepest slope of all variants across all dilutions, being significantly different than BA.1, BA.4 and BA.5 at some dilutions, though never reached significance compared to WA1/2020 due to a slightly higher standard error. BA.1 had the least steep slope at all dilutions. At the two highest viral inputs, 1e5 and 1e4 TCID_50_, WA1/2020 had a steeper slope than BA.1, though the trend did not continue for the lower dilutions (Fig. 2B).

**Fig. 2.**
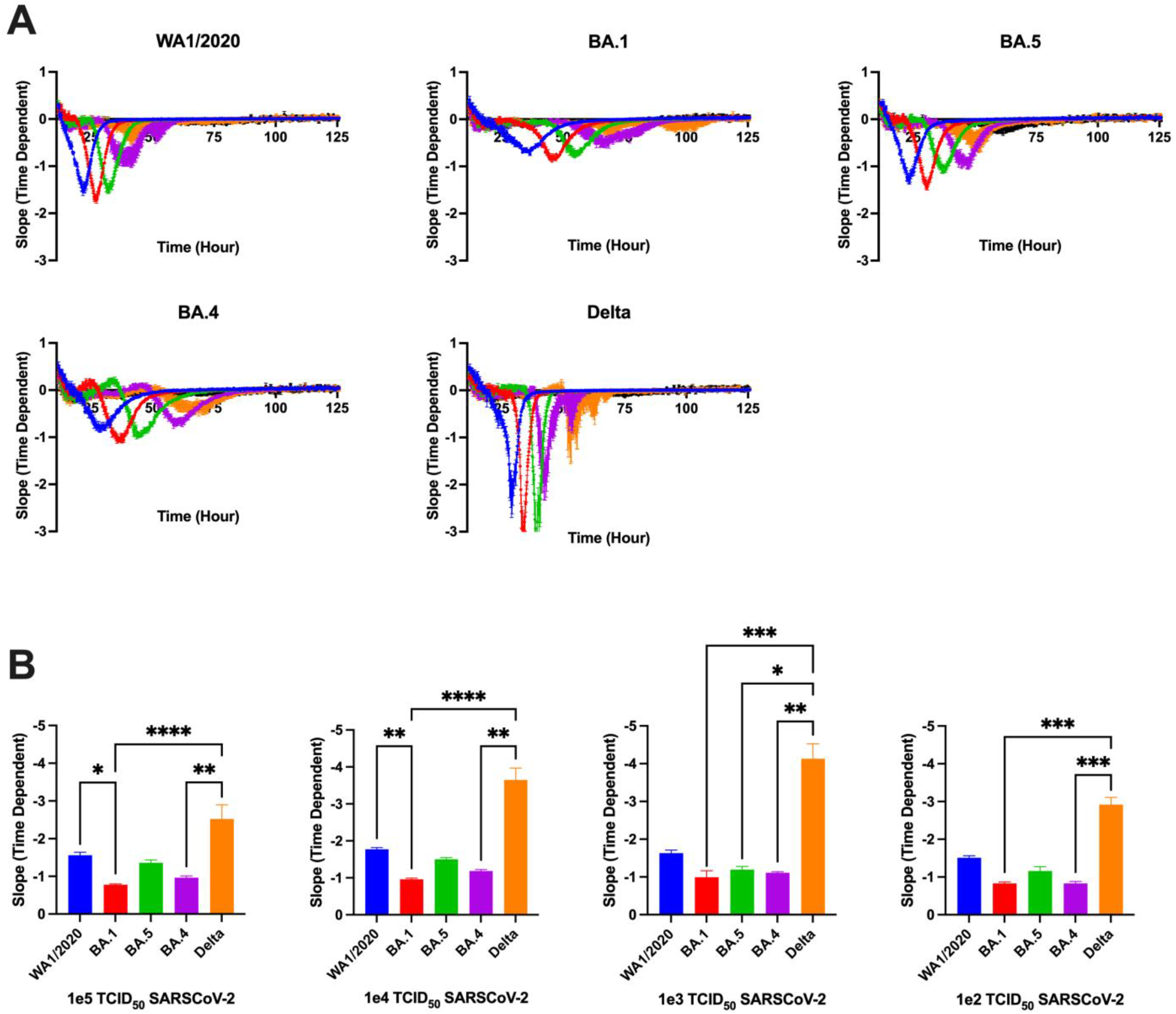
Cell Index over time and slope of Vero/TMPRSS2 monolayers inoculated with SARS-CoV-2 Variants. Cell index was determined every 15s over the course of 5 days. Data was normalized at 11.756h. (**A**) The time taken to reach each variant’s max slope across multiple viral concentrations graphed as averages with SEM and **(B)** Value of max slope. Groups were compared via Kruskal-Wallis test. (*, p<0.05; **, p<0.01; ***, p<0.001; ****, p<0.0001).

### AUC relationships correlate with slope characteristics and concentration of viral inoculum

Comparisons of slope characteristics reveal relationships between various aspects of the cell index curves. The input virus concentration does not appear to correspond with the value of max slope, except for Delta at 1e3 to 1e5 TCID_50_ dilutions. Relationships do exist between concentration and time to max slope, as well as concentration and AUC, with both time and AUC falling as viral input concentration increases (Fig. 3A).

**Fig. 3.**
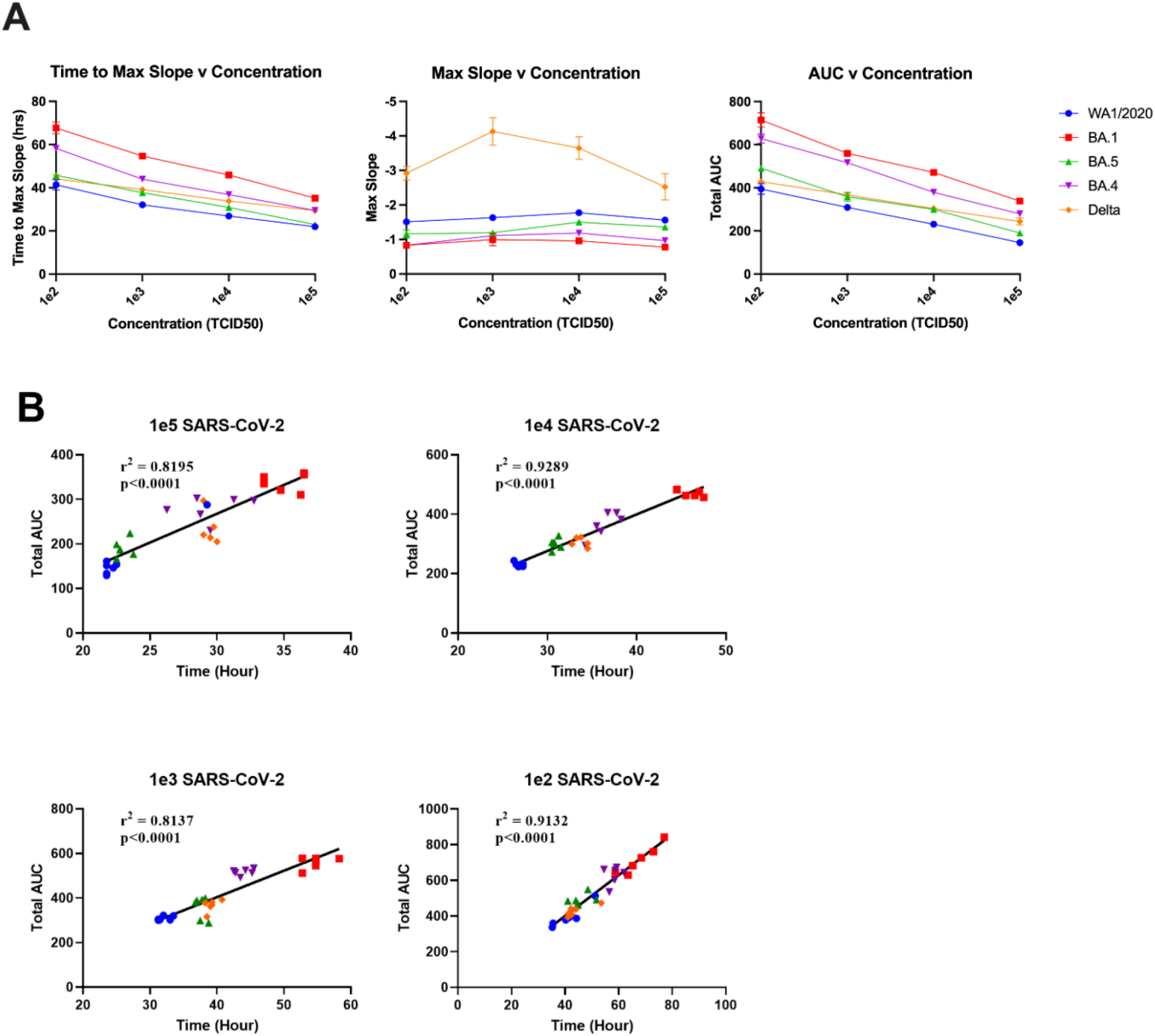
Slope and AUC Relationships of SARS-CoV-2 Variants. (A) Relationships between Time to max slop, value of max slope, and AUC with varying viral concentrations applied to cell monolayers. (B) Relationship between AUC and time to reach max slope for each variant. P value represents Spearman correlation.

Time to max slope does correlate with AUC across all dilutions, with 1e4 and 1e2 TCID_50_ input concentrations having the highest correlations, at 0.9289 and 0.9132, respectively. All p values were below 0.0001, indicating a high degree of correlation (Fig. 3B). The AUC and value of the max slope are fairly related, with clustering occurring among variants. The exception to this is Delta, with only small clustering at the highest viral input, and very little among other dilutions (Fig. S2). Time to max slope and value of max slope are somewhat similar overall to the AUC/max slope value relationship. Variants other than Delta cluster together, giving the impression of some degree of interrelatedness (Fig. S3).

### Trends in slope characteristics follow visually captured cell death

Images of each monolayer at the same spot in each well were captured at 60-minute intervals over the course of the experimental period allowing for generation of time lapse videos. This enables cytopathic effect (CPE) to be interpreted visually during viral infection with each variant. At 1e5 TCID_50_ input, WA1/2020 reaching widespread cytotoxicity can be seen before other variants, with BA.1 showing the slowest progression. The progression of Delta can be seen as a slow initial development of CPE, followed by a rapid progression across the monolayer once CPE begins. Overall, Delta is most similar to BA.4 and BA.5 visually, with WA1/2020 and BA.1 being the most rapid and most slow, respectively. This trend mirrors that seen in the slope characteristics (Fig. S4). Comparisons between variants taken at the same point, 5 hours prior to the max slope timepoint, show a contrast between Delta and other variants. At concentrations of 1e4 and 1e5, the effect of cellular fusion appears evident from visual inspection of the monolayer (Fig. 4). The effect is less apparent but still visible at lower concentration (Fig. S5).

**Fig. 4.**
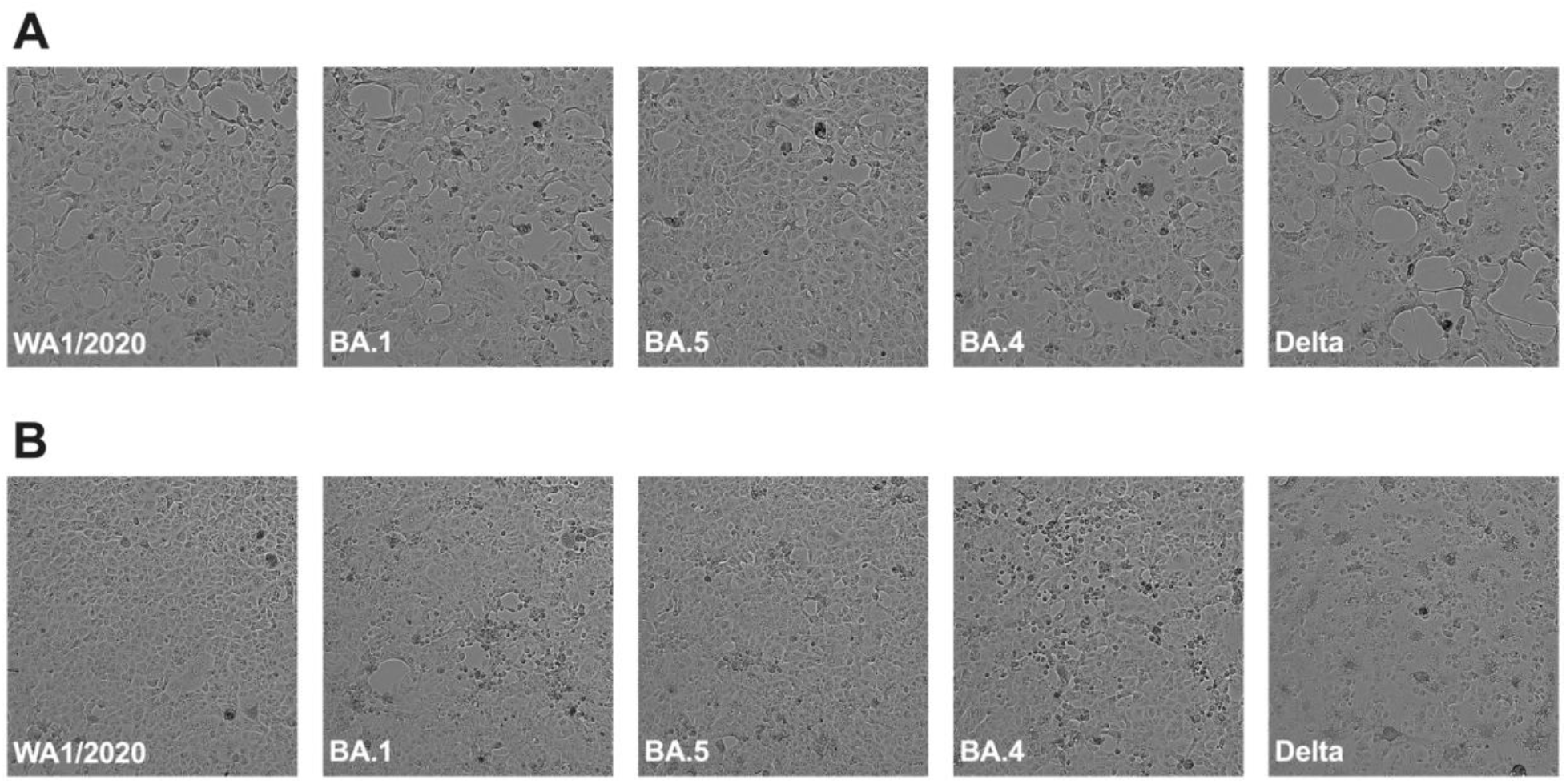
Monolayers visualized during replication. Images of monolayers taken 5 hours prior to each variant’s maximum slope time point for Vero/TMPRSS2 inoculated with (A) 1e5 TCID_50_ or (B) 1e4 TCID_50_ SARS-CoV-2.

## Discussion

Here we investigated the differences in the replication kinetics of multiple SARS-CoV-2 variants of concern via the surrogate cell index method that measures electrical impedance of a cell monolayer. This correlates to monolayer integrity that is altered by viral replication and cytopathic effect. We utilized this technique to quantify multiple aspects of the cell index curves generated over a time course of monolayer infection with known viral input concentrations, including AUC, time to max slope, and value of max slope.

Cell index patterns over time as monolayer infection progresses can be seen as a decrease in impedance, or the ability of the monolayer to resist current. The patterns here, such as the slope of that decrease, can be characterized in order to determine differences in viral kinetics of replication. WA1/2020 reached max slope more quickly than other variants, indicating that it reaches a point during replication of cellular destruction on a wide scale more quickly than others. BA.5 was the closest to this rapid pace, and BA.1 was the slowest. These data suggest that mutations accumulated or lost in BA.5 relative to BA.1 explain this kinetic difference and may indicate selection to regain a greater level of replicative ability lost in the BA.1 variant. Whether these changes have any impact on pathogenicity is difficult to address. Indeed, if any of the newly emerging subvariants of Omicron have recapitulated the pathogenicity of earlier, pre-Omicron variants, this effect might be masked in humans by the high global incidence of prior vaccination or infection, particularly with the BA.1 variant, which has led to very high levels of global immunity, which undoubtedly reduces disease and death associated with subsequent infection with any variant. Thus, any increase in pathogenicity associated with specific variants should be rigorously addressed in animal models.

The absolute value of the slope correlates to how quickly the monolayer is losing its ability to resist current, indicating viral destruction of the monolayer. Delta was the highest variant, by far, for this metric, indicating that it rapidly destroys the monolayer, despite its delay reaching that slope as compared to WA1/2020. As before, BA.1 was the slowest here, indicating a slower replication potential. It is intriguing to speculate what might explain Delta’s uniqueness in this measure. Other studies have indicated that Delta replicates faster than other variants using different measures than we used here [41] but those measures may fail to identify the nuanced difference between Delta and the ancestral WA1/2020 that RTCA captures in our assays. When Delta was initially detected, there was much research into identifying mutations that might have led to its increased replicative capacity and possibly pathogenicity. One mutation in particular, spike P681R, was shown to increase both fusogenicity and pathogenicity in hamsters [42,43]. Whether this mutation underlies the effects we have identified remains to be seen but could be tested.

Total area under the curve, indicating the total amount of time of replication in the monolayer, was the highest for BA.1, reinforcing the slow replication profile. WA1/2020 was the most rapid overall, with BA.5 being closely followed by Delta, indicating Delta doesn’t match WA1/2020 despite its rapid pace of monolayer destruction once initiated.

Our data reveal or confirm several aspects of viral replication that may shed light on the ongoing COVID-19 pandemic. First, we found that the Omicon subvariant BA.5 recapitulates most replicative features of the ancestral WA1/2020 variant, suggesting selection to regain features missing in the BA.1 variant. Second, our data reveal a unique replication profile for the Delta variant, which has been shown to induce greater pathogenicity in animal models and possibly in humans as well. Finally, and perhaps most importantly, our data add to a large body of work demonstrating that the original Omicron variant, now termed BA.1, was demonstrably and significantly less fit in terms of replicative ability than any other tested variant, which very well may correlate with the clearly reduced pathogenicity of this variant. That this variant swept the globe, infecting far more people than all other variants, likely suggests that its immense immune evasion capacity far outweighed its reduced replicative ability allowing for widespread infection including in those with pre-existing immunity.

Together, the work described here further elucidates the patterns of replication exhibited by each variant of SARS-CoV-2, with added clarity of real time cell analysis allowing us greater insight into potential replication kinetics across time points not typically examined. Real time cell analysis is a robust method that, in conjunction with established tools including qPCR, genomics, animal modeling and public health surveillance, will give us greater insight into the unique nature of newly emerging variants.

