## Supplemental Figures for "Real time analysis of SARS-CoV-2 induced cytolysis reveals distinct variant-specific replication profiles"

**Fig. S1. Mean Cell Index over time of Vero//TMPRSS2 monolayers inoculated with SARS-CoV-2 Variants.**

The absolute value of each variant's max slope across multiple viral concentrations represented as means without error. Legend indicates quantity of virus in $TCID_{50}$ added onto cell monolayers.

**Fig. S2. Comparisons of AUC and Max Slope Value**

Relationship between AUC and absolute value of max slope for each variant.

**Fig. S3. Comparisons of Max Slope Value and Time to Max Slope**

Relationship between absolute value of max slope and time to max slope for each variant at different $TCID_{50}$ values.

**Fig. S4. Comparison videos**

Time-synched videos representative of CPE induced by each SARS-CoV-2 variant on Vero/TMPRSS2 monolayers.

**Fig. S5. Monolayers visualized during replication**

Images of monolayers taken 5 hours prior to each variant's maximum slope time point for Vero/TMPRSS2 inoculated with (A) 1e3 $TCID_{50}$ or (B) 1e2 $TCID_{50}$ SARS-CoV-2.

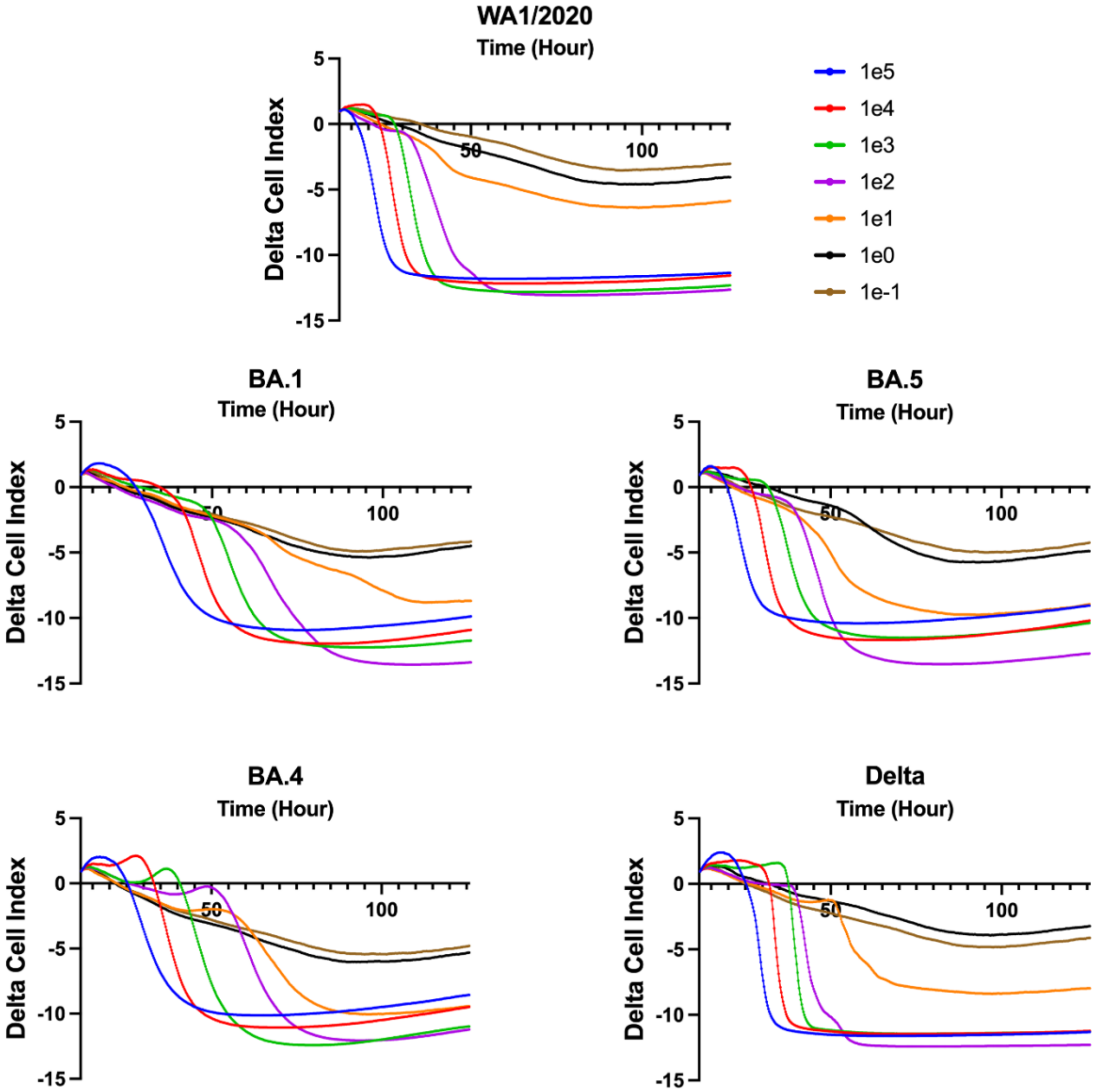
WA1/2020
Time (Hour)
Delta Cell Index
1e5
1e4
1e3
1e2
1e1
1e0
1e-1
BA.1
BA.5
BA.4
Delta
5
0
-5
-10
-15
50
100

**Fig. S1.**

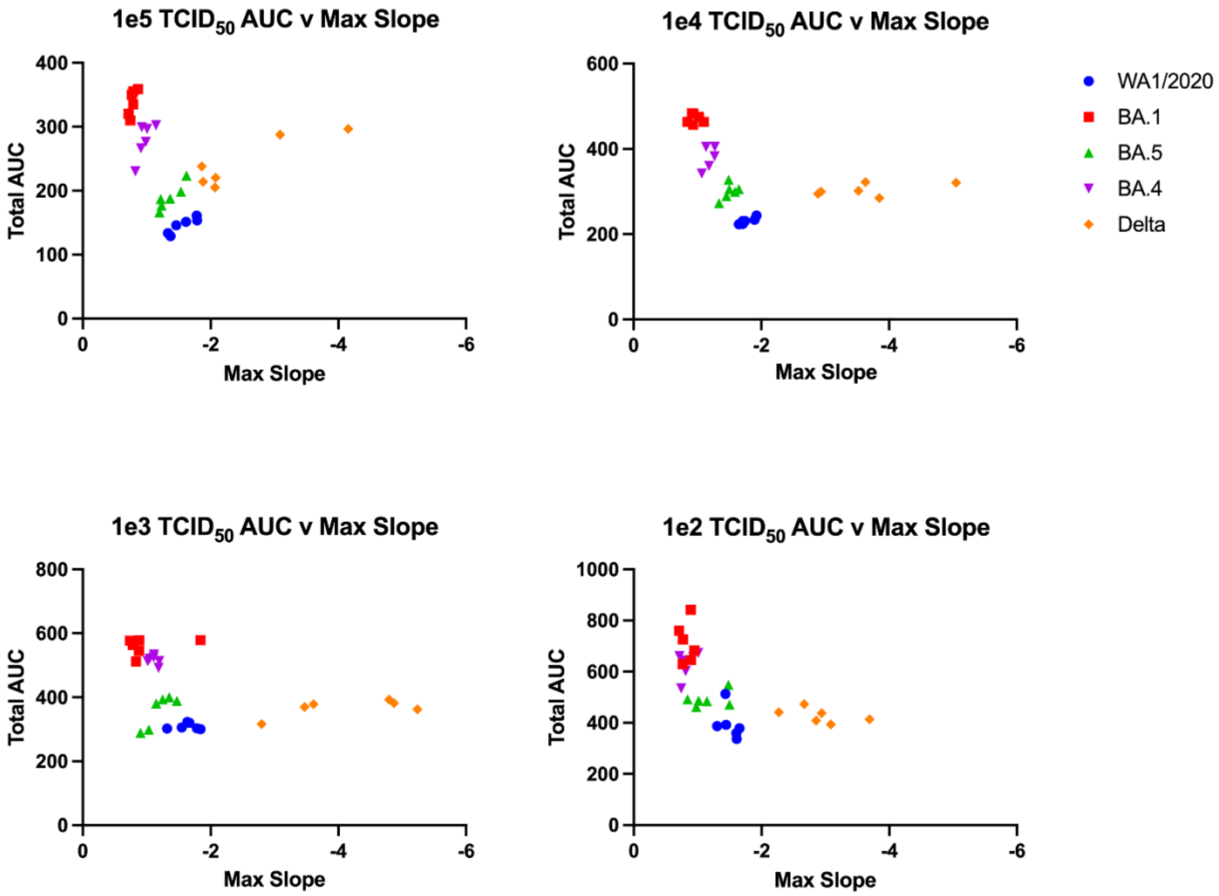

1e5 TCID50 AUC v Max Slope
1e4 TCID50 AUC v Max Slope
1e3 TCID50 AUC v Max Slope
1e2 TCID50 AUC v Max Slope
Total AUC
Max Slope
WA1/2020
BA.1
BA.5
BA.4
Delta


**Fig. S2.**

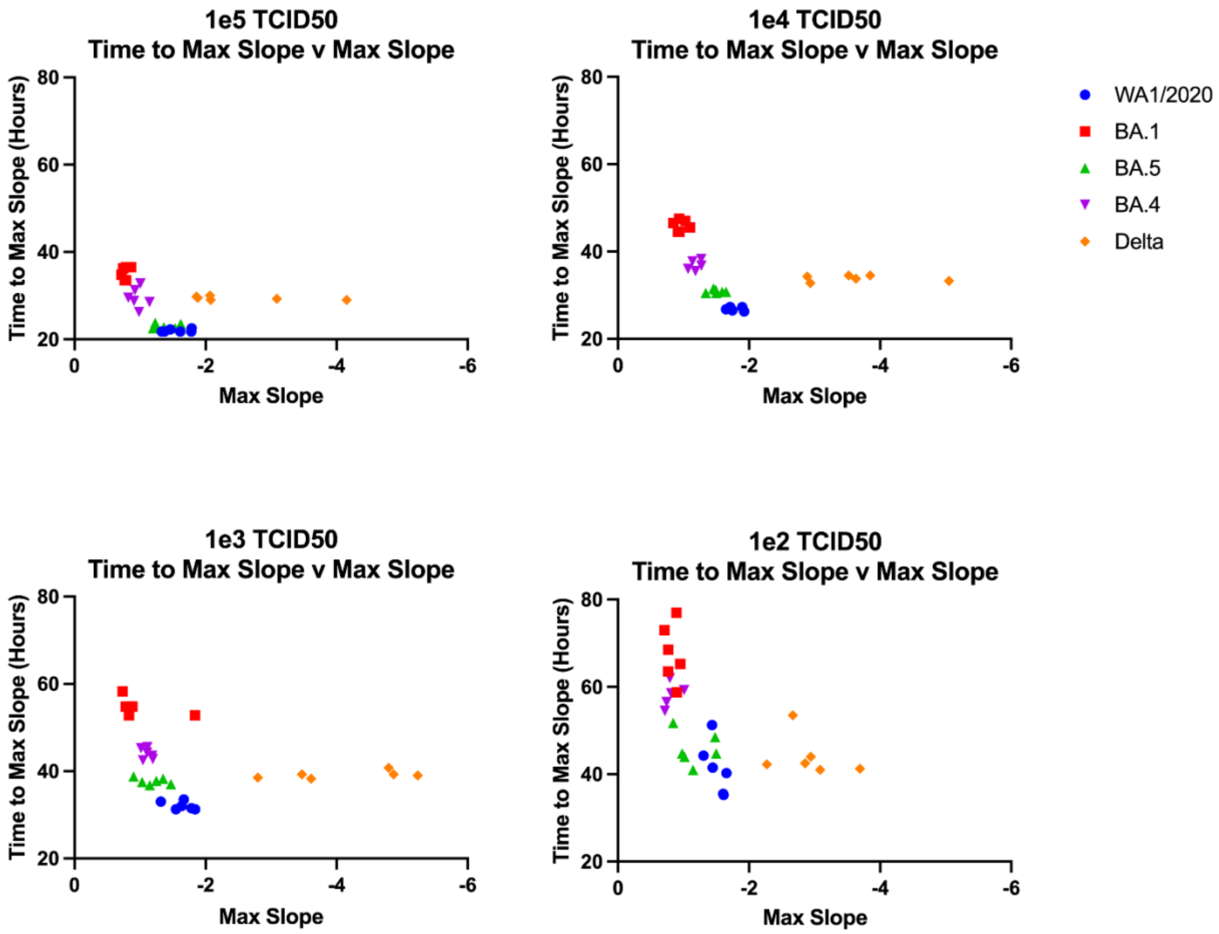
1e5 TCID50
Time to Max Slope v Max Slope
Time to Max Slope (Hours)
Max Slope
1e4 TCID50
Time to Max Slope v Max Slope
Time to Max Slope (Hours)
Max Slope
1e3 TCID50
Time to Max Slope v Max Slope
Time to Max Slope (Hours)
Max Slope
1e2 TCID50
Time to Max Slope v Max Slope
Time to Max Slope (Hours)
Max Slope
WA1/2020
BA.1
BA.5
BA.4
Delta

**Fig. S3.**

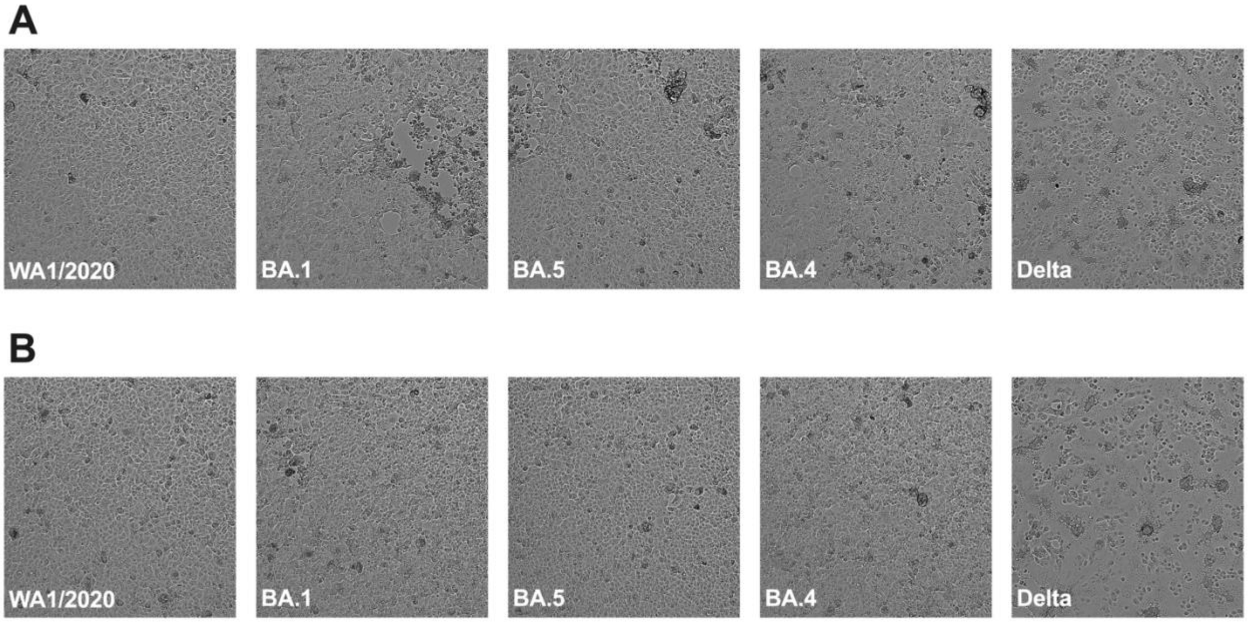


**Fig. S5.**
